# Small-molecule inhibitors of the RNA m^6^A demethylase FTO potently support the survival of dopamine neurons

**DOI:** 10.1101/2021.02.23.432419

**Authors:** Simona Selberg, Li-Ying Yu, Olesja Bondarenko, Esko Kankuri, Neinar Seli, Vera Kovaleva, Koit Herodes, Mart Saarma, Mati Karelson

## Abstract

The fat mass and obesity-associated protein (FTO), an RNA N^6^-methyladenosine (m^6^A) demethylase, is an important regulator of central nervous system development, neuronal signalling and disease. We present here the target-tailored development and biological characterization of small-molecule inhibitors of FTO. The active compounds were identified using high-throughput molecular docking and molecular dynamics screening of the ZINC compound library. In FTO binding and activity-inhibition assays the two best inhibitors demonstrated K_d_ = 185 nM; IC_50_ = 1.46 μM (compound **2**) and K_d_ = 337 nM; IC_50_ = 28.9 μM (compound **3**). Importantly, the treatment of mouse midbrain dopamine neurons with the compounds promoted cellular survival and rescued them from growth factor deprivation induced apoptosis already at nanomolar concentrations. Moreover, these inhibitors demonstrated good blood-brain-barrier penetration in the model system, 31.7% and 30.8%, respectively. The compounds **2** and **3** protected dopamine neurons with greater potency than our recently developed alkylation repair homolog protein 5 (AlkBH5) m^6^A demethylase inhibitors. Inhibition of m^6^A RNA demethylation by small-molecule drugs, as presented here, has therapeutic potential and provides tools for the identification of disease-modifying m^6^A RNAs in neurogenesis and neuroregeneration. Further refinement of the lead compounds identified in this study, can also lead to unprecedented breakthroughs in the treatment of neurodegenerative diseases.

## Introduction

Chemical modifications of RNA have a critical impact on many cellular functions, such as proliferation, survival and differentiation^1,2^. In eukaryotic messenger RNA, the most abundant modification is N^6^-methyladenosine (m^6^A), which affects RNA splicing, intracellular transport, translation, and cytoplasmic degradation of RNA^3,4^. The levels of m^6^A in RNA are regulated by specific enzymes, methyltransferases, demethylases and m^6^A reader proteins. These include m^6^A writers such as the methyltransferase-like protein 16 (METTL16)^5^ as well as the RNA methyltransferase enzyme complex METTL3/METTL14/WTAP usually composed of three components: METTL3 (methyltransferase-like 3), METTL14 (methyltransferase-like 14) and WTAP (Wilm’s tumour-1-associated protein)^6,7^. RNA m^6^A eraser enzymes include the RNA demethylases FTO (fat mass and obesity-associated protein)^8,9^ and AlkBH5 (alkylation repair homolog protein 5)^10^. Additionally, the fate of RNA in post-transcriptional processes is determined by the m^6^A reader proteins that recognize specific m^6^A-modified RNA sequences and affect the stability, translation and/or cellular localization of the transcript. Several RNA reader proteins have been identified^7,11^, including three members of YTH N^6^-Methyladenosine RNA Binding Protein (YTHDF)-family (YTHDF 1-3) and two members of the YTH domain-containing protein (YTHDC)-family (YTHDC1-2)^12^. Collectively, these three types of proteins coordinate the m^6^A RNA methylome and its fate in the eukaryotic cells.

RNA m^6^A modifications have been assigned key orchestrating roles in brain development, neuronal signalling and neurological disorders^13–18^. For example, m^6^A-dependent mRNA decay has been shown to be critical for proper transcriptional pre-patterning in mammalian cortical neurogenesis^19^. Weng et al. demonstrated that axonal injury-induced m^6^A methylation and downstream signalling enhances the synthesis of regeneration-associated proteins essential for functional axon regeneration of peripheral sensory neurons^20^. Moreover, it has been shown that genes associated with m^6^A control may play a role in conferring risk of dementia^21^. However, because the homeostasis of RNA m^6^A methylation in neurons is controlled on multiple levels, unselective or global modification of m^6^A levels can yield contradictory results. For example, through the actions of the reader protein YTHDF1, m^6^A residues have been shown to directly facilitate adaptive processes such as learning and memory in the adult mouse hippocampus^22^. Deficiency of the m^6^A eraser FTO has been demonstrated to lead to impaired learning and memory through reduced proliferation and neuronal differentiation of adult neural stem cells in *Fto* full-knockout mice^23^. In addition, conventional and dopamine (DA) neuron-specific *Fto* gene knockout mice show impaired DA receptor type 2 (D2R)- and type 3 (D3R)-dependent control of neuronal activity and behavioral responses^24^. Treatment of PC12 cells derived from a pheochromocytoma of the rat adrenal medulla *in vitro* as well as the rat striatum *in vivo* with the 6-OHDA neurotoxin in a rat model of Parkinson’s disease (PD) results in a global reduction of the m^6^A residues in mRNAs^25^. Reduction of m^6^A levels in pheochromocytoma PC12 cells by treatment with the non-selective nucleoside methylation inhibitor, cycloleucine, or alternatively by FTO overexpression induced apoptotic cell death through increased expression of N-methyl-D-aspartate (NMDA) receptor 1, oxidative stress and Ca^2+^ influx^25^. However, these results do not demonstrate directly the dysregulation of m^6^A in dopamine neurons. Nevertheless, because the striatum contains the fibres of DA neurons, it is possible that the dysregulation of m^6^A is related to neurodegeneration in PD. Available data indicate that either the compensatory upregulation or downregulation of m^6^A could be needed in neuronal cells, depending on their physiological or pathological state. However, the precise role of the RNA demethylases FTO and AlkBH5 in the regulation of survival and regeneration of DA neurons has remained enigmatic. One reason for this is the lack of highly specific inhibitors of these enzymes. As shown in the rat 6-OHDA PD model, the downregulation of m^6^A in the striatum occurs in parallel with the axonal degeneration and DA neuron death^25^.Therefore it is logical to hypothesize that inhibitors of the RNA m^6^A demethylases FTO or AlkBH5 should support the homeostasis of m^6^A in DA neurons and their survival under stress. Only a very limited number of FTO inhibitors are presently known, mostly of non-specific nature^26–31^.

We used *in silico*-based rational target-tailored development of small-molecule FTO inhibitors and determined their binding affinity, kinetics, and their effect on enzymatic functions experimentally. We identified unique small-molecule ligands that bind to FTO and very potently inhibit its enzymatic activity. In particular, two of these FTO inhibitors, already at 10 nM concentration, supported the survival of growth factor-deprived primary DA neurons in culture. Two AlkBH5 inhibitors that we have described earlier^32^, were less potent in rescuing DA neurons. This is the first demonstration that inhibitors of FTO, and in general m^6^A regulators can support the survival and protect dopamine neurons from growth factor deprivation induced death *in vitro.* These compounds may further serve as lead molecules for development of novel drugs for neurodegenerative diseases such as PD.

## Methods

### Compounds

4-aminoquinoline-3-carboxylic acid (**1**); Vitas-M Laboratory, Ltd, Cat. No. STK660776, Purity: >90%.

4-amino-8-chloroquinoline-3-carboxylic acid (**2**); Vitas-M Laboratory, Ltd, Cat. No. STK787835, Purity: >90%.

8-aminoquinoline-3-carboxylic acid (**3**); Ark Pharm, Ltd, Cat. No. AK200350, Purity: 98%.

(2E)-4-[(3-methylphenyl)formohydrazido]-4-oxobut-2-enoic acid (**4**); Vitas-M Laboratory, Ltd, Cat. No. STK120795, Purity: >90%.

8-hydroxyquinoline-5-carboxylic acid (**5**); Enamine, Ltd, Cat. No. Z233564176, Purity: >90%.

3-methyl-N’-(3-methylbenzoyl)benzohydrazide (**6**); Vitas-M Laboratory, Ltd, Cat. No. STK087016, Purity: >90%.

2-[(1-hydroxy-2-oxo-2-phenylethyl)sulfanyl]acetic acid (**7**); Enamine Ltd., Cat No. EN300-14040, Purity: >90%.

4-{[(furan-2-yl)methyl]amino}-1,2-diazinane-3,6-dione (**8**); Vitas-M Laboratory, Ltd., Cat. No. STL352808, Purity: >90%.

### FTO protein

FTO protein was provided by ProteoGenics SAS (Schiltigheim, France, www.ProteoGenix.science.com). The details of the protein synthesis are given in Supplementary material. The purity of the FTO protein was >90%.

### Computational modeling

The crystal structure of the FTO in complex with 5-carboxy-8-hydroxyquinoline IOX1 (pdb:4IE4)^27^ was chosen for the prediction of potential efficient ligands using molecular docking modeling. The ligand IOX1 was removed from the complex in order to proceed with the search of novel ligands. The catalytic center of the protein involves a bivalent transition metal ion, either Mn^2+^, Fe^2+^, Ni^2+^ or Zn^2+^. In our molecular docking simulations Zn^2+^ was used. The raw crystal structure was corrected and hydrogen atoms were automatically added to the protein using Schrödinger’s Protein Preparation Wizard of Maestro 10.7^33^. AutoDock 4.2^34^ was used for the docking studies to find out binding modes and binding energies of ligands to the receptor. The number of rotatable bonds of ligand was set by default by AutoDock Tools 1.5.6^34^. However, if the number was greater than 6, then some of rotatable bonds were made as non-rotatable, otherwise calculations can be inaccurate. The active site was surrounded with a grid-box sized 80 × 80 × 80 points with spacing of 0.375 Å. The AutoDock 4.2 force field was used in all molecular docking simulations. The docking efficiencies (DE) were calculated as follows

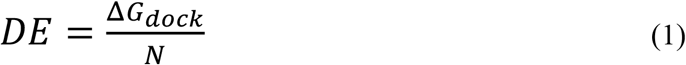

where Δ*G*_*dock*_ is the docking free energy and N – the number of non-hydrogen (“heavy”) atoms in the ligand molecule.

The structures of ligand molecules were optimized using the density functional theory B3LYP method^35^ with 6-31G basis set.

The molecular dynamics simulations were carried out using Desmond simulation package of Schrödinger LLC^36^. The NPT ensemble with the temperature 300 K and pressure 1 bar was applied in all runs. Five simulation runs with the length 10 ns and with relaxation time 1 ps were carried out for each system. The OPLS_2005 force field parameters were used in all simulations^37^. The long-range electrostatic interactions were calculated using the particle mesh Ewald method^38^. The cutoff radius in Coulomb’ interactions was 9.0 Å. The water molecules were described using the simple point charge (SPC) model^39^. The types of physical interactions between the ligands and enzyme were analyzed using the Simulation Interaction Diagram tool implemented in Desmond molecular dynamics package. The stability of molecular dynamics simulations was monitored by looking on the root mean square deviation (RMSD) of the ligand and protein atom positions in time.

### Protein binding study using microscale thermophoresis

The microscale thermophoresis (MST) experiments were performed using Monolith NT.115 instrument (NanoTemper Technologies GmbH, Germany). Recombinant human FTO protein was labeled with His-tag using Monolith His-Tag Labeling Kit RED-tris-NTA (NanoTemper Technologies GmbH; MO-L008). The labelled FTO protein (target) was used at 20 nM in all the experiments and 10 μM starting concentrations of ligands FTO inhibitors **2** or **3** were used in both series of experiments.

The measurements were done in a 10mM Na-phosphate buffer, pH 7.4, containing 1 mM MgCl_2_, 3 mM KCl, 150 mM NaCl, 0.05% Tween-20 in premium coated capillaries (NanoTemper Technologies GmbH; MO-K025) using red LED source, power set at 100% and medium MST power at 25°C. Each data point represents mean fraction bound values from n=3 independent experiments per binding pair ±S.D, K_d_ values ± error estimations are indicated. Data analysis was performed using MO.Affinity Analysis v2.3 software (NanoTemper Technologies GmbH).

### Enzyme inhibition assay

The enzymatic assay was modified from Huang et al.^29^ The experiments were conducted in reaction buffer (50 mM Tris-HCl, pH 7.5, 300 μM 2OG, 280 μM (NH_4_)_2_Fe(SO_4_)_2_ and 2 mM L-ascorbic acid). The reaction mixture contained 200 ng methylated N^6^-adenine RNA probe (SEQ ID NO: 1) (5’-CUUGUCAm^6^ACAGCAGA-3’, Dharmacon, Lafayette, CO) and 10 nM FTO protein and different concentrations of ligands (1 nM to 100 μM). Reactions were incubated on 96-well plate for 2 h at RT. After that, the amount of m^6^A that was measured using EpiQuik m^6^A RNA methylation Quantification Colorimetric Kit (Epigentek, Farmingdale, NY).

The inhibitory effect *IE* of compounds on RNA probe demethylation by FTO was calculated as the enhancement of the m^6^A amount as compared to the negative control (DMSO) relative to the difference between m^6^A amounts of the positive control (max inhibition) and the negative control (eq. (2)):

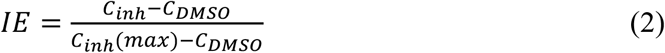

where *C_inh_, C_inh_(max) and C_DMSO_* are the amounts of m^6^A at a given concentration of the inhibitor, maximum inhibition and in the case of DMSO, respectively.

### Primary cultures of midbrain dopamine neurons and m^6^A regulator treatment

The midbrain floors were dissected from the ventral mesencephali of 13 days old NMRI strain mouse embryos following the published procedure^40^. The tissues were incubated with 0.5% trypsin (103139, MP Biomedicals, Inc, Thermo Fisher Scientific, Waltham, MA) in HBSS (Ca^2+^/Mg^2+^-free) (14170112, Invitrogen, Thermo Fisher Scientific, Carlsbad, CA) for 20 min at 37°C, then mechanically dissociated. Cells were plated onto the 96-well plates coated with poly-L-ornithine (Sigma-Aldrich, Merck KGaA, St. Louis, MO). Equal volumes of cell suspension were plated onto the center of the dish. The DA neurons were cultured for 5 DIV in presence of cell culture media Dulbecco’s MEM/Nut mix F12 (Invitrogen/Gibco; 21331– 020), 1xN2 serum supplement (Invitrogen/ Gibco; 17502–048), 33mM D-Glucose (Sigma; G-8769), 0.5 mM L-Glutamine (Invitrogen/ Gibco; 25030–032), and 100 μg/ml Primocin (Invivo Gen, San Diego, CA)]. On 6 DIV, to deprive growth factors, the cultures were washed three times with normal medium and the FTO and AlkBH5 inhibitors or glial cell line-derived neurotrophic factor (GDNF) were applied^40^. The cells were grown for 5 days with different concentrations of FTO and AlkBH5 inhibitors. Human recombinant GDNF (100 ng/ml) (Icosagen AS, Tartu, Estonia) or a condition without any neurotrophic compound added were used as positive and negative controls, respectively.

After growing 5 days, the neuronal cultures were fixed and stained with anti-Tyrosine Hydroxylase antibody (MAB318, Millipore Bioscience Research Reagents, Temecula, CA). Images were acquired by CellInsight (Thermo Fisher Scientific) high-content imaging equipment. Immunopositive neurons were counted by CellProfiler software and the data was analyzed by CellProfiler analyst software^41^. The results are expressed as % of cell survival compared to GDNF-maintained neurons^42^.

### Artificial blood-brain barrier model

Artificial *in vitro* blood-brain barrier (BBB) was established as described by Le Joncour et al.^43^ Murine endothelial cells bEnd3 (ATCC CRL-2299) were co-cultured with murine hypoxia-inducible factor knock out (HIFko) astrocytes (passage 12, received from Le Joncour) on hanging cell culture inserts (BD Falcon 353091, Franklin Lakes, NJ) for 5 days in fetal bovine serum (FBS)-free conditions. 2.5 ml of FTO inhibitors (final concentration of 10 μM) were added into the insert (representing the “blood” side) of the BBB cell. After 1 h incubation, 1 ml of the sample was taken from the insert and from the well (“brain” side), and the concentration of the compounds was measured by high-performance liquid chromatography (HPLC). Penetration % was defined by dividing the concentration of the compound in the well compared to the concentration in the insert. The concentrations of the studied compounds is each side of the BBB cell were analyzed using LC-MS system consisting of Agilent 1290 UHPLC and Agilent 6460 Triple Quadrupole MS (both from Agilent Technologies Inc, Santa Clara, CA). Chromatographic separation was carried out on Zorbax RRHD Eclipse Plus (3 × 100 mm, 1.8 μm) column using 0.1% aqueous formic acid and acetonitrile as eluent components. Linear gradient from 3% to 100% acetonitrile in 6 minutes followed by 3 min isocratic segment. Eluent flow rate was 0.5 ml/min and sample injection volume was 1 μl. Electrospray ion source (Agilent Jet Stream) was operated in positive ionization mode using default values for gas flows, temperatures and potentials. For each compound two transitions were selected and respective collision energies coarsely optimized.

### Quantification and statistical analysis

Enzymatic assay curve-fitting analysis and determination of the IC_50_ and EC_50_ values were performed using AAT Bioquest, Inc. Quest Graph™ IC_50_ Calculator (v.1, Sunnyvale, CA). MST data analysis was performed using MO.Affinity Analysis v2.3 software (NanoTemper Technologies GmbH, Munich, Germany). Statistical significance in cell survival experiments was assessed using one-way ANOVA and unpaired t test with the GraphPad Prism8 software. Results were considered statistically significant at p values lower than 0.05.

## Results and Discussion

### Computational modeling of FTO ligand binding site and virtual screening

The regions of probable interactions between a ligand and FTO protein were found by carrying out the molecular docking using AutoDock 4.1. As shown by the molecular docking calculations, the amino acid residues of the protein Asp233, Tyr106, Glu234, Arg96 and Arg322 were involved in specific interactions between the protein and ligand (Figure 1)

**Figure 1.**
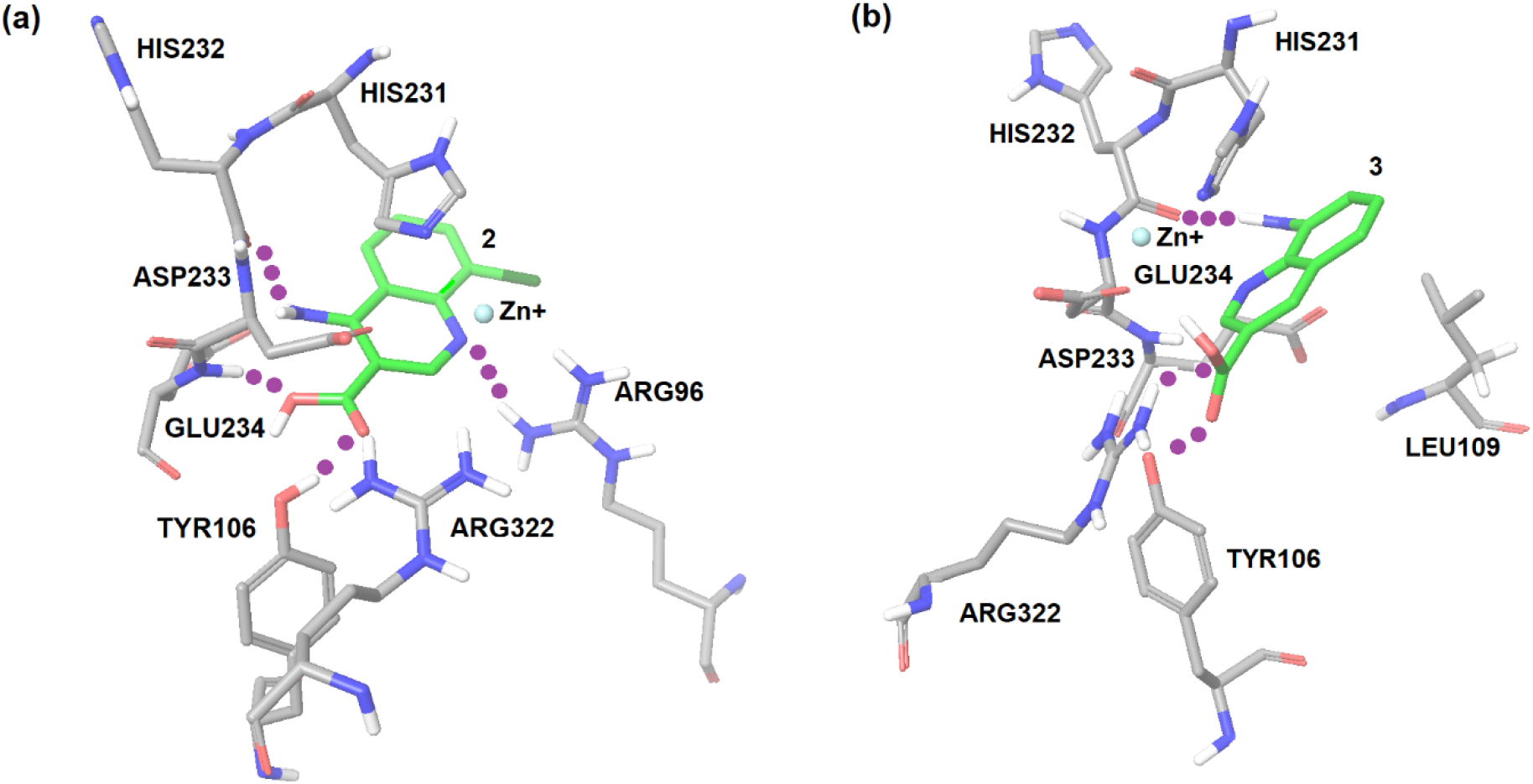
Docking modeling FTO binding sites of the compounds **2** (a) and **3** (b).

A virtual screening on ZINC compound library^44^ was carried out using the best known FTO inhibitors from the ChemBL database^45^ as templates (Figure 2). The docking free energies ΔG and ligand efficiencies LE of the best binding compounds are given and their molecular structures are given in Table 1.

**Figure 2.**
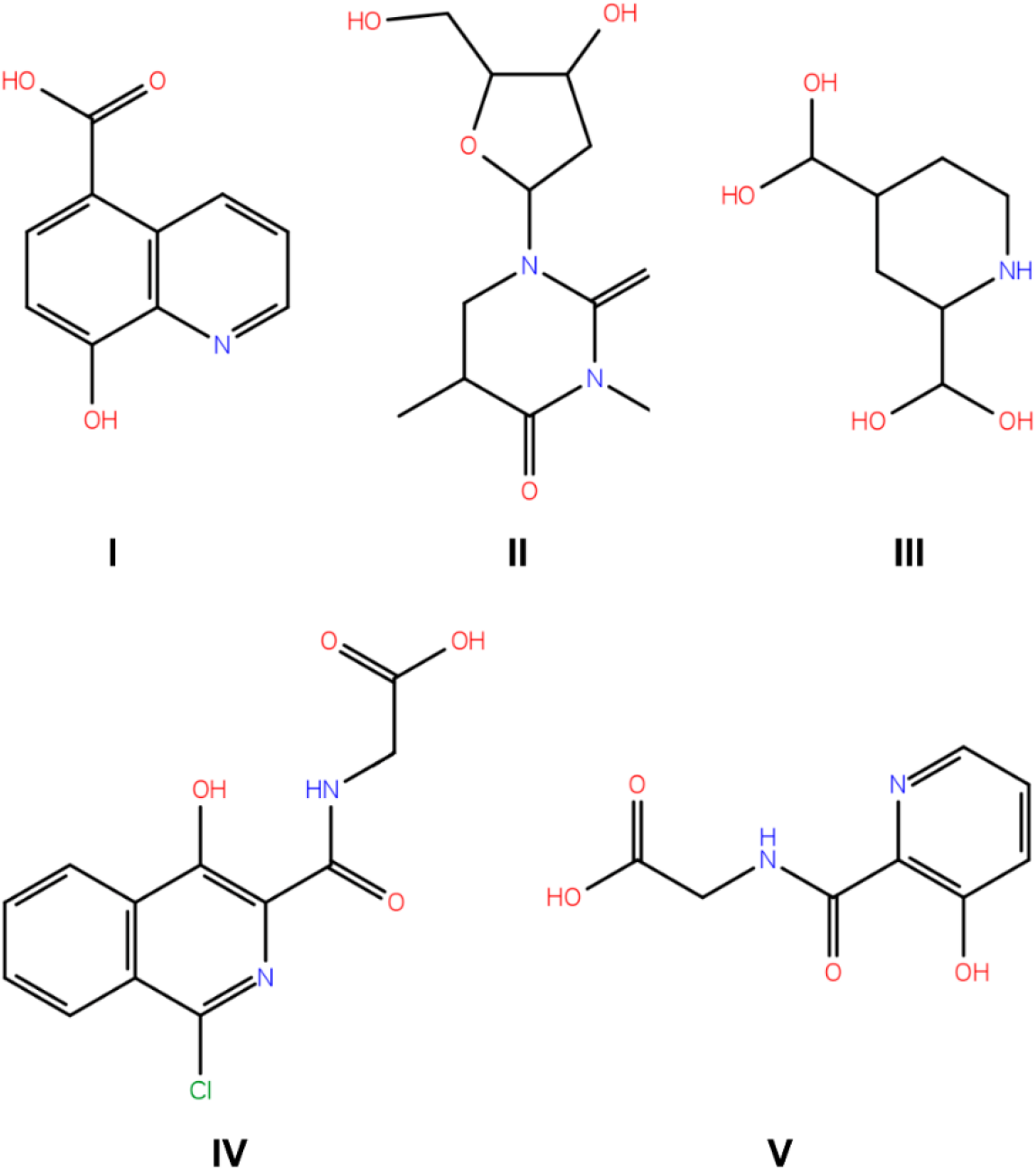
A set of the known FTO inhibitors used as templates in high-throughput virtual docking screening.

**Table 1.**
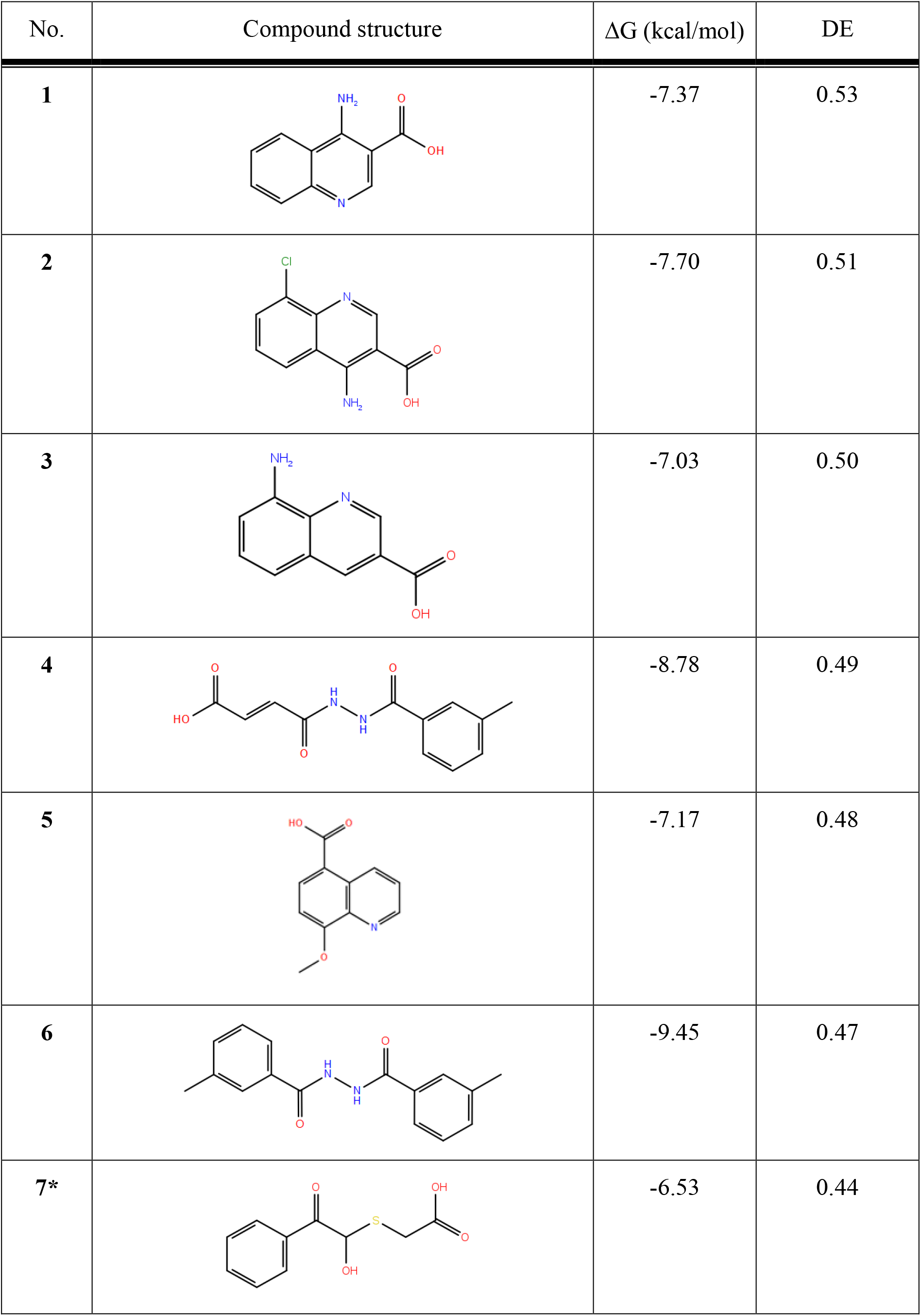
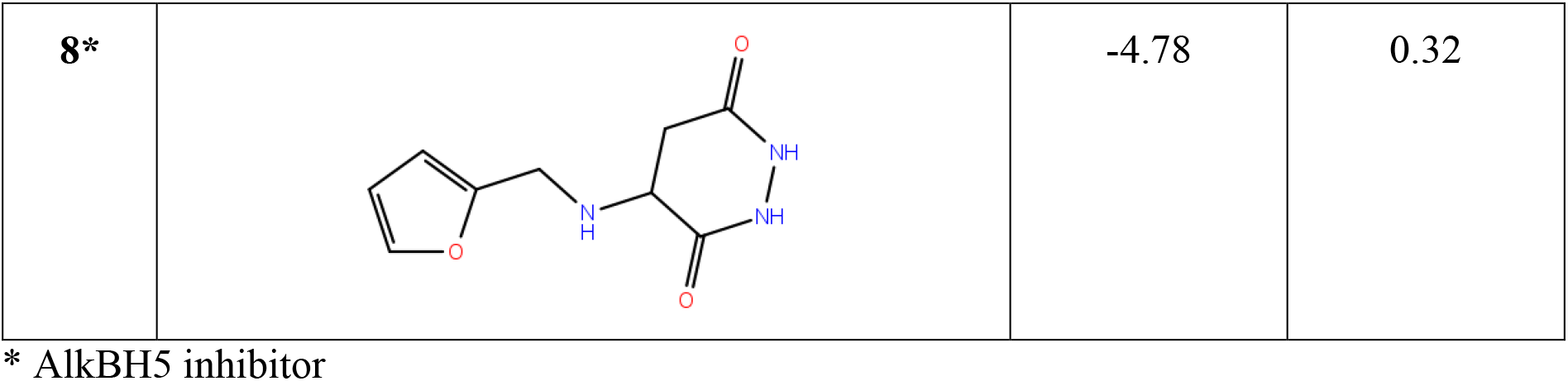
The compounds with the highest docking efficiencies DE to FTO protein.

| No. | Compound structure | $\Delta G$ (kcal/mol) | DE |
| --- | --- | --- | --- |
| 1 |  | -7.37 | 0.53 |
| 2 |  | -7.70 | 0.51 |
| 3 |  | -7.03 | 0.50 |
| 4 |  | -8.78 | 0.49 |
| 5 |  | -7.17 | 0.48 |
| 6 |  | -9.45 | 0.47 |
| 7* |  | -6.53 | 0.44 |
| 8* |  | -4.78 | 0.32 |

\* AlkBH5 inhibitor

The molecular dynamics simulations were carried out for two compounds, the compounds with the best enzymatic inhibition activity (**2** and **3**).

In the case of compound **2**, the several molecular dynamics simulation runs were carried out with the length of 10 ns. This system was stable throughout the calculation time (Figure 3a). A very strong hydrogen bond is detected between the pyridine nitrogen atom of the ligand and the ammonium group of Arg96 residue of the FTO protein (Figure 3b). The simulation interactions diagram (Figure 3c) indicates that the most important interactions for this compound are hydrogen bonds between ligand and residues Arg96, Glu234, Arg322 and Asp233 of FTO. Furthermore, there are additional hydrophobic interactions between ligand **2** and FTO protein. The bars in diagram Figure 3c characterize the time fraction that a particular specific interaction is maintained during the simulation. Based on this, we can assume that the compound **2** is bound to tight specific pocket at the active site of FTO protein (Figure 3d)^46^.

**Figure 3.**
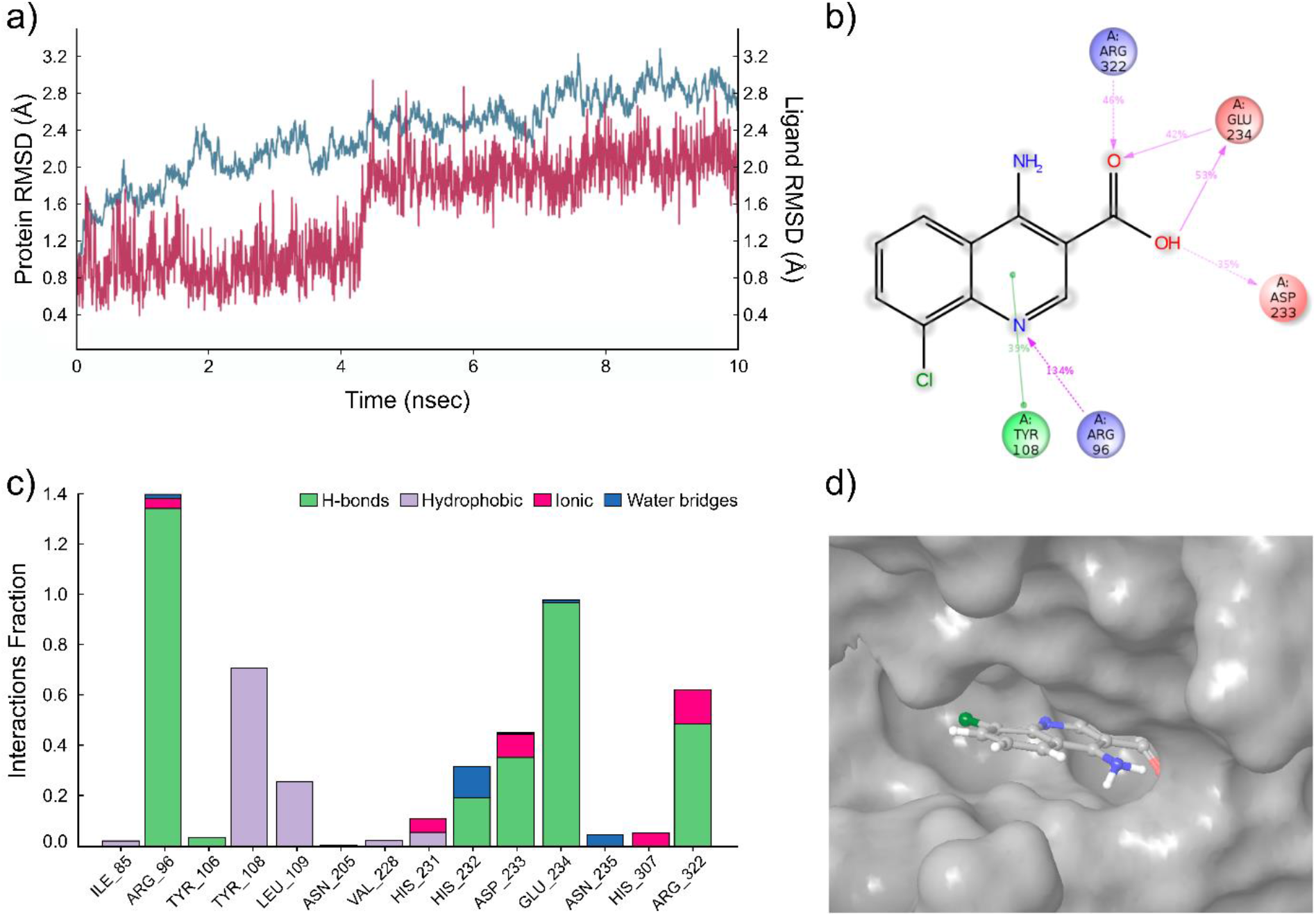
The results of the molecular dynamics simulation of the FTO complex with compound **2**. (a) The protein and ligand position root mean square deviation (RMSD) plot against time in the case of the FTO complex with compound **2** for a representative 10 ns run. (b) Desmond 2D profile data for the compound **2** binding to FTO protein. (c) Normalized stacked bar chart representation of interactions and contacts over the course of trajectory (values over 1.0 are possible as some residue make multiple contacts of same subtype with ligands); interactions occurring more than 50% of the simulation time. Interaction diagram between the compound **2** and FTO protein. (d) The position of the compound **2** in the structure of FTO, related to Figure 1a.

The results of the molecular dynamics simulation of compound **3** are summarized in Figure 4. Again, five molecular dynamics simulation runs were carried out with the length of 10 ns, and the trajectory analysis shows the stability of the system during the calculation (Figure 4a). The results indicate the presence of hydrogen bonds between the ligand carbonyl group of compound **3** and Glu234 and Asp233 of the FTO protein. In addition, a water bridge with Arg96 and salt bridge with Arg322 (Figure 4b) is suggested. The simulation interactions diagram (Figure 4c) reveals a very stable hydrogen bonding (Asp233 and Glu234) and several ionic bridges (His231, Asp233, His307 and Arg322) and water bridges (Arg96, Ser229 and Arg322) between the compound **3** and protein. The compound is bound to tight specific pocket at the active site of FTO protein (Figure 4d).

**Figure 4.**
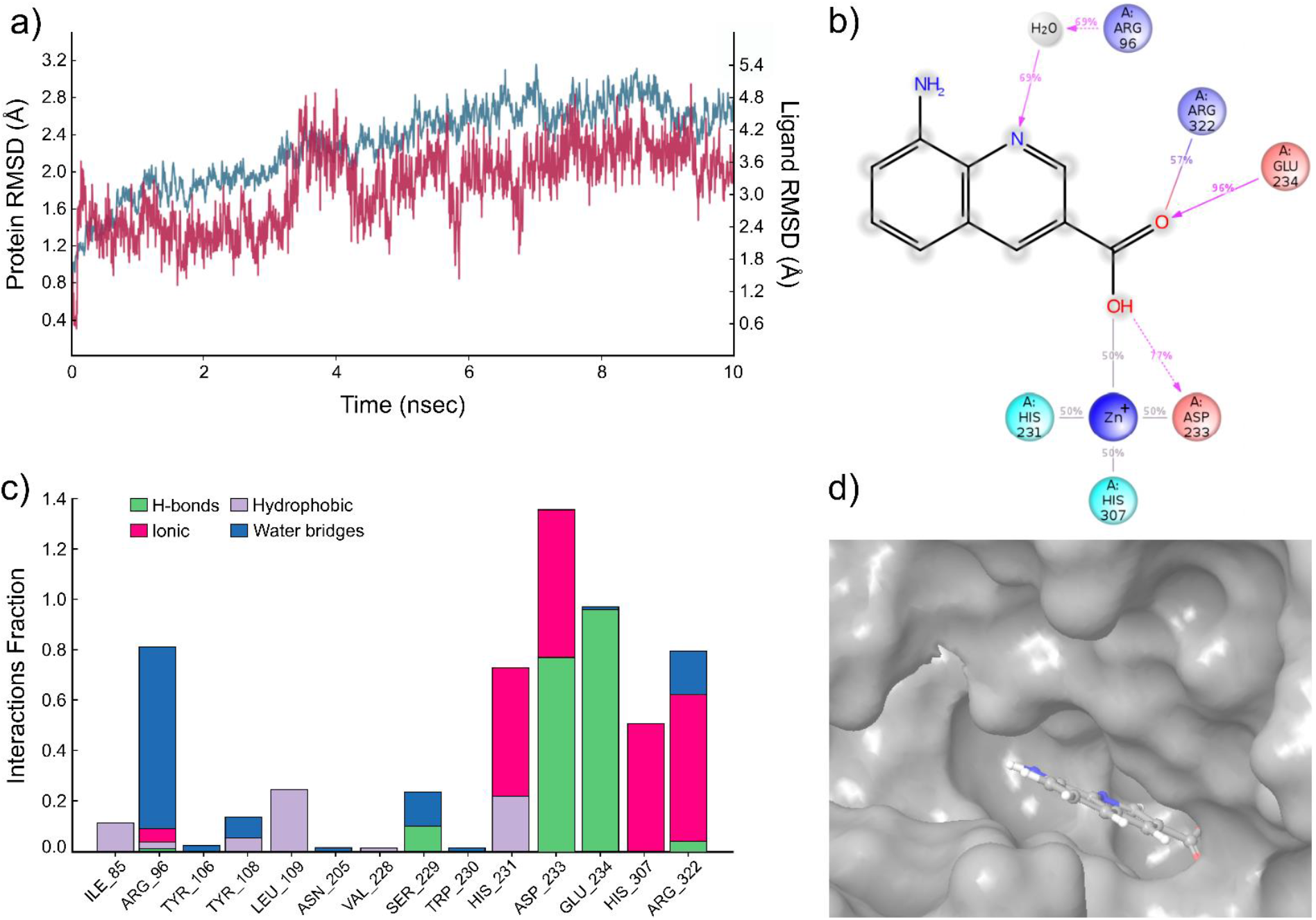
The results of the molecular dynamics simulation of the FTO complex with compound **3**. (a) The protein and ligand position root mean square deviation (RMSD) plot against time in the case of the FTO complex with compound **3** for a representative 10 ns run. (b) Desmond 2D profile data for the compound **3** binding to FTO protein. (c) Normalized stacked bar chart representation of interactions and contacts over the course of trajectory (values over 1.0 are possible as some residue make multiple contacts of same subtype with ligands); interactions occurring more than 50% of the simulation time. Interaction diagram between the compound **3** and FTO protein. (d) The position of the compound **3** in the structure of FTO, related to Figure 1b.

### Enzyme activity inhibition

The enzyme inhibition measurements were carried out for the predicted FTO strongly bound compounds **1** – **6**. A significant concentration-dependent inhibitory effect was observed for quinolone derivatives **2** and **3** (Figure 5). The inhibitory concentrations were IC_50_ = 1.46 μM for compound **2** and IC_50_ = 28.9 μM for compound **3**. No significant inhibitory effect was noticed for the other three predicted compounds (**1**, **4**, **5** and **6**) up to the 100 μM concentration.

**Figure 5.**
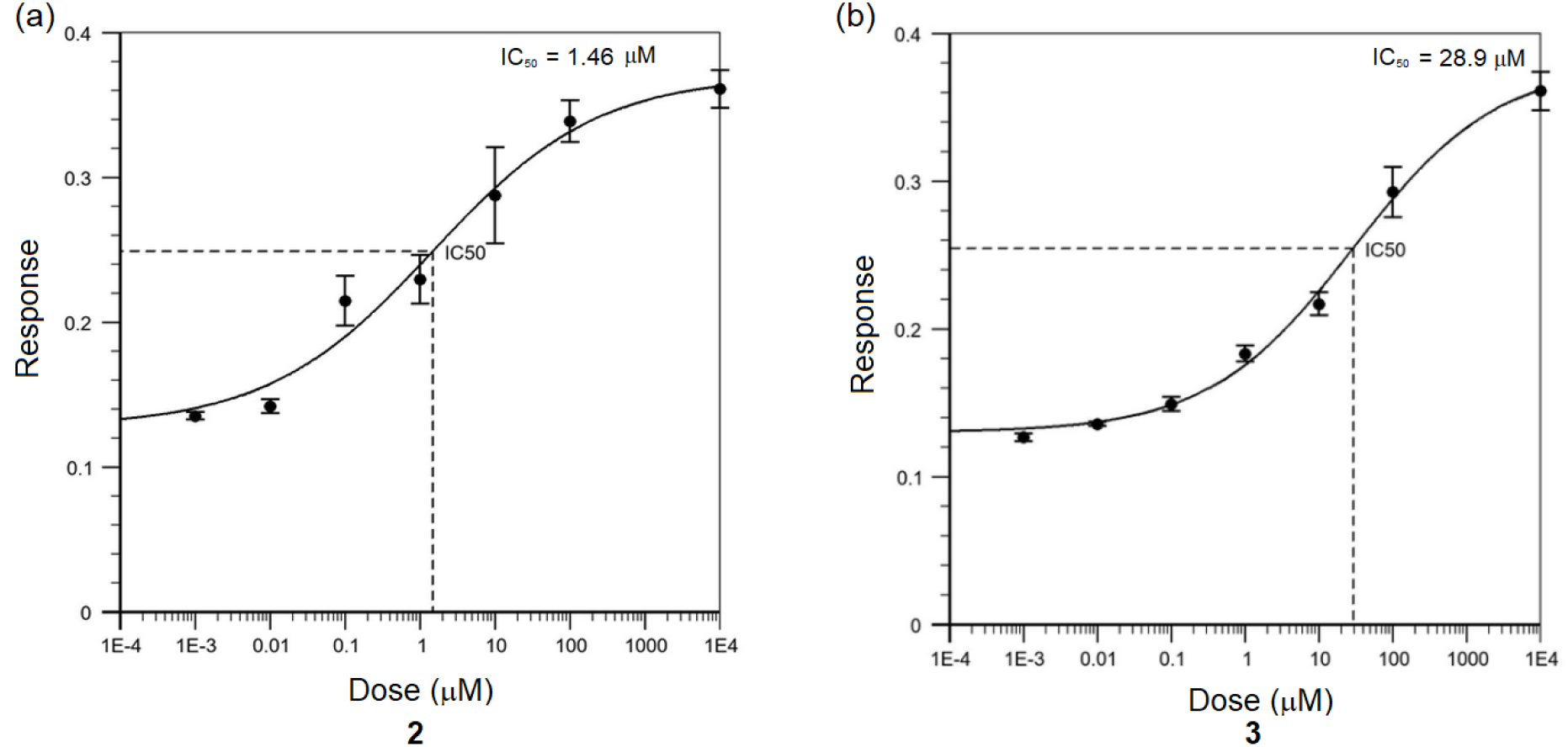
Inhibitory effects of the compounds **2**. (a) and **3** (b) on the demethylation of the probe RNA by FTO. Response indicates the m^6^A level in probe.

### Protein binding of compounds

Furthermore, we studied the binding of the two active inhibitors **2** and **3** to the FTO protein using MST. Both compounds are binding at sub-micromolar concentrations. The protein binding K_d_ values K_d_ = 185 ± 77 nM for compound **2** and K_d_ = 337 ± 184 nM for compound **3** (Figure 6) are in good agreement with the respective enzymatic inhibition IC_50_ values for these compounds.

**Figure 6.**
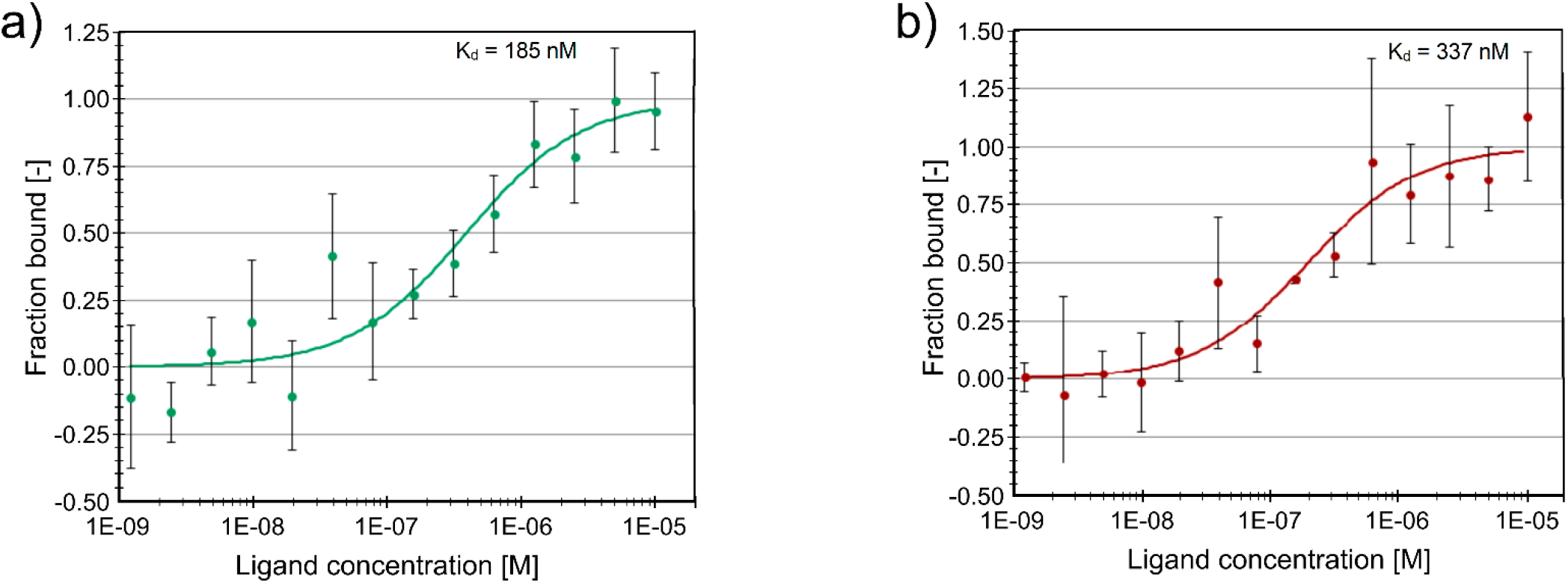
Two inhibitors directly interact with FTO protein as shown by microscale thermophoresis. (a) Unlabeled titrated compound **2** (0-10 μM) interacts with Alexa647-labeled through His-tag FTO (20 nM). (b) Unlabeled titrated compound **3** (0-10 μM) interacts with Alexa647-labeled through His-tag FTO (20 nM). For (a) and (b) microscale thermophoresis binding curves, showing fraction bound values from n=3 individual repeats per binding pair ±SEM, K_d_ values±error estimations are indicated.

### Neuronal survival experiments

To our knowledge, the direct effect of FTO and AlkBH5 inhibitors has never been tested on DA neurons. Earlier data demonstrate that reduced m^6^A levels in 6-OHDA-treated tyrosine hydroxylase-expressing rat pheochromocytoma PC12 cells having some similarity to peripheral sympathetic neurons by overexpressing FTO result in apoptosis^25^. We therefore hypothesized that the inhibition of RNA m^6^A demethylases in DA neurons could counteract this apoptotic process. To test this hypothesis, we carried out a study on the influence of the developed FTO inhibitors on the survival of mouse midbrain dopamine neurons after inducing their apoptosis by growth factor deprivation. It has been demonstrated that the preferred cellular substrate for FTO is not the m^6^A but its further modification N^6^2’-O-dimethyladenosine (m^6^Am), which is exclusively found adjacent to the 7-methylguanine (m^7^G) cap in mRNA^47–49^. Furthermore, FTO is primarily, and potentially exclusively localized in the nucleus^8,50^. Thus, the presently known main m^6^A demethylating enzyme is the AlkBH5, belonging to the non-heme Fe^(II-)^ and α-KG-dependent dioxygenase AlkB family of proteins. Contrasting FTO, AlkBH5 has no activity towards m^6^Am and appears to be localized to nuclear speckles^9^. There may be also a significant difference in the target RNAs. In the case of FTO, the main target RNA may not be mRNA, but snRNA^51^.

Since mRNA and snRNA m^6^A modification may have impact on neuronal survival, it was therefore interesting to compare the effects of inhibition of these two RNA m^6^A demethylases on the survival of dopamine neurons in the *in vitro* model of Parkinson’s disease. Thus, the experiments were carried out not only with two FTO inhibitors developed in this study, compounds **2** and **3**, but also with two our recently reported AlkBH5 inhibitors, 2-[(1-hydroxy-2-oxo-2-phenylethyl)sulfanyl]acetic acid **7** and 4-{[(furan-2-yl)methyl]amino}-1,2-diazinane-3,6-dione **8**. The enzyme inhibitory concentrations against AlkBH5 are IC_50_ = 0.840 μM for the compound **7** and IC_50_ = 1.79 μM for the compound **8**, respectively^32^.

GDNF has been shown to protect cultured embryonic dopamine neurons from growth factor deprivation induced, as well as 6-OHDA-induced cell death *in vitro* and *in vivo*^40,52^. We, therefore, assessed the neuroprotective ability of different concentrations of FTO or AlkBH5 inhibitors in cultured growth factor deprived dopamine neurons. Human recombinant GDNF (100 ng/ml) or a condition without any neurotrophic compound added were used as positive and negative controls, respectively. Growth factor deprivation caused cell death by 50-70 %. The results expressed as % of cell survival compared to GDNF-maintained neurons for the FTO inhibitors are presented in Figures 7a and 7b. Both FTO inhibitors **2** and **3** similarly to GDNF dose-dependently protected embryonic midbrain dopamine neurons in culture from growth factor deprivation -induced cell death. A neuroprotective effect can be seen already at 10 nM, and statistically significant outcome is observed at the concentration 100 nM and 1000 nM concentrations of both the inhibitors **2** and **3**. Hence, the inhibition of the m^6^A demethylase FTO promotes on the survival of dopamine neurons and rescues them in growth factor deprivation *in vitro* model of Parkinson’s disease without any signs of toxicity of the tested compounds.

**Figure 7.**
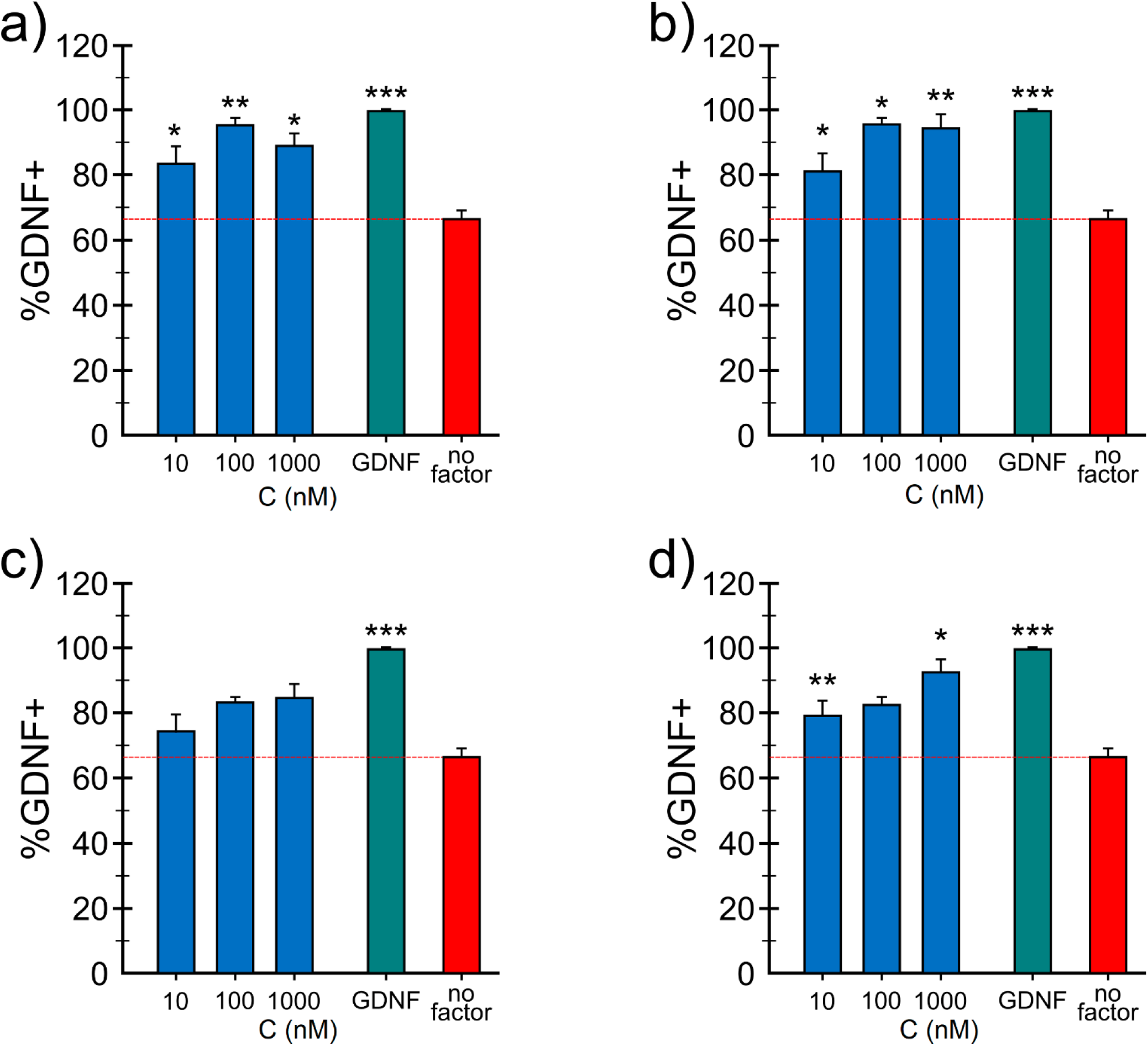
The effect of the FTO inhibitors **2** (a) and **3** (b) and AlkBH5 inhibitors **7** (c) and **8** (d) on the survival of the dopamine neurons. The number of TH-positive cells in the wild-type midbrain cultures with growth factor deprivation and treated with vehicle, FTO inhibitors **2** (a) and **3** (b) and AlkBH5 inhibitors **7** (c) and **8** (d) or GDNF normalized to the total number of cells in the culture and presented as percentage of vehicle treated samples, average from 4-5 well from one experiment. Both experiments were repeated 3 times with reproducible results. Concentration of GDNF used as a positive control is 3 nM (100 ng/ml) and concentration of inhibitors are in nM. No factor - Vehicle. * P < 0.05, ** P < 0.01, *** P < 0.001, one-way ANOVA with Dunnett’s *posthoc* test.

Similarly, in growth factor deprivation model, AlkBH5 inhibitors **7** and **8** at three tested concentrations increased the number of TH-positive neurons (Figures 7c and 7d, respectively). Compound **8** rescued growth factor deprivation challenged dopamine neurons at 10 nM and at 1000 nM, but showed only a trend at 100 nM. Compound **7** on the other hand showed only trend in the neuroprotection of apoptosis induced E13 dopamine neurons.

It is interesting to note that the potency of FTO inhibitors **2** and **3** in protecting and rescuing DA neurons *in vitro* is comparable to that of GDNF. Since GDNF, when directly injected into the midbrain, protects dopamine neurons also in animal models of PD,^52,53^ it is logical to assume that FTO and AlkBH5 inhibitors can also be neuroprotective *in vivo*. The main limitation in the clinical use of GDNF and other neurotrophic proteins in the treatment of PD is their inability to pass through the BBB. As the first step towards *in vivo* testing of the neuroprotective activities of FTO inhibitors we assessed their ability to penetrate through the artificial BBB.

### Penetration of the FTO inhibitors through the blood-brain barrier in artificial in vitro model

The effects of FTO inhibitors were studied in the artificial *in vitro* BBB model, where murine endothelial cells bEnd3 were co-cultured with murine HIFko astrocytes on hanging cell culture inserts. The BBB penetration % defined as the ratio of the concentration of the compound in the well (“brain”) side and the concentration in the insert, were 31.7 ±3.3 % for compound **2** and 30.8±1.9% for compound **3**. Therefore, both compounds exhibit good BBB penetration ability.

## Conclusions

The m^6^A RNA modifications and their dynamics in the cell have been recently related to numerous cell developmental, physiological and pathological processes, including neurogenesis and neuronal survival. Here we demonstrated that the inhibition of the m^6^A demethylation by inhibiting FTO or AlkBH5, that supposedly takes place predominantly in the cell nucleus supports the survival of the dopamine neurons and protects them from growth factor deprivation -induced apoptosis. The neuroprotective efficacy of two FTO inhibitors in this *in vitro* model of PD is similar to that of GDNF. Since both neuroprotective FTO inhibitors have the ability to pass through the artificial BBB *in vitro*, it is of great interest to test their activity in animal models of PD in future studies. Differently from GDNF, these compounds have the potential to penetrate the BBB and therefore they can be potentially delivered systemically avoiding risky and complicated brain surgery. It has been demonstrated earlier that the substrate RNA targets for the two RNA m^6^A demethylases, FTO and AlkBH5 are different, m^6^Am and m^6^A, respectively. Therefore, it is exciting to observe a very similar (although of different intensity) effects by the inhibitors of both these enzymes on the dopamine neuron survival. The dopamine neurons have complicated neurite network with extensive number of synaptic contacts that require very demanding intracellular RNA transport and mRNA translation. Further studies in order to identify these specific RNA targets in neuronal cells could give new basic information about the neurogenesis and neuroregeneration. Another important outcome of the present study is the demonstration of possibility to employ a completely new type of neuroprotective compounds, the small-molecule RNA m^6^A demethylase inhibitors for the further development of drugs against Parkinson’s and possibly Alzheimer’s diseases, amyotrophic lateral sclerosis and other neurodegenerative disorders.

## Author contributions

S.S. computational modeling, virtual screening, enzyme inhibition experiments, data analysis; L.Y. neuronal survival experiments; O.B. and K.H. BBB measurements; V.K. MST binding experiments; M.S. design and data analysis of neuronal survival and MST experiments; N.S. funding acquisition and data analysis; M.K. conceptualization and coordination of the study, data analysis; M.K., M.S., S.S., E.K., V.K. writing and editing manuscript.

