## Supplementary material for "Small-molecule inhibitors of the RNA m^6^A demethylase FTO potently support the survival of dopamine neurons"

### *Protein synthesis*

(1) Gene coding for FTO protein.

The cDNA coding for the target proteins were chemically synthesized with optimization for expression in mammalian cells. A Kozak sequence was added in 5' position, and a sequence coding for a 6His-tag was added in 3' position of the protein. The respective sequence is given below.

***>FTO-6His cDNA - 1542 bp***

```
ATGAAGAGAACACCCACCGCCGAGGAGAGGGAGAGAGAGGCCAAGAAGCTGAGGCTGCTGGAGGAGCTGGAGGATACATG
GCTGCCTTACCTGACACCCAAGGATGATGAGTTCTACCAGCAGTGGCAGCTGAAGTACCCCAAGCTGATCCTGAGGGAGG
CCTCCAGCGTGAGCGAGGAGCTGCACAAGGAGGTGCAGGAGGCCTTCCTGACCCTGCACAAGCACGGCTGTCTGTTTAGA
GACCTGGTGAGGATCCAGGGCAAGGACCTGCTGACCCCCGTGAGCAGAATCCTGATCGGCAATCCCGGCTGTACCTACAA
GTACCTGAACACCAGGCTGTTCACCGTGCCTTGGCCCGTGAAGGGCAGCAACATCAAGCACACCGAGGCCGAGATCGCCG
CCGCCTGTGAGACCTTTCTGAAGCTGAACGATTACCTGCAGATCGAGACAATCCAGGCCCTGGAGGAGCTCGCCGCCAAG
GAGAAGGCCAATGAGGACGCCGTGCCCCTGTGCATGAGCGCCGACTTCCCCAGGGTGGGCATGGGCAGCTCCTACAATGG
CCAGGATGAGGTGGACATCAAGTCCAGAGCCGCCTACAACGTGACACTGCTGAACTTTATGGATCCCCAGAAGATGCCCT
ACCTGAAGGAGGAGCCTTACTTTGGCATGGGCAAGATGGCCGTGTCCTGGCACCACGATGAGAATCTGGTGGACAGGTCC
GCCGTGGCCGTGTACTCCTACAGCTGCGAGGGCCCTGAGGAGGAGTCCGAGGATGATAGCCACCTGGAGGGCAGGGACCC
CGACATCTGGCACGTGGGCTTTAAGATCAGCTGGGACATCGAGACACCTGGCCTGGCCATCCCCCTGCACCAGGGAGATT
GTTACTTTATGCTGGACGATCTGAACGCCACACACCAGCACTGTGTGCTGGCCGGCTCCCAGCCCAGATTCTCCTCCACA
CACAGAGTGGCCGAGTGCTCCACCGGCACCCTGGATTACATCCTGCAGAGATGCCAGCTGGCCCTGCAGAATGTGTGCGA
TGATGTGGATAATGATGATGTGTCCCTGAAGAGCTTCGAGCCCGCCGTGCTGAAGCAGGGCGAGGAGATCCACAATGAGG
TGGAGTTCGAGTGGCTGAGACAGTTCTGGTTCCAGGGCAACAGATACAGGAAGTGCACAGACTGGTGGTGTCAGCCCATG
GCCCAGCTGGAGGCCCTGTGGAAGAAGATGGAGGGCGTGACCAATGCCGTGCTGCACGAGGTGAAGAGGGAGGGCCTGCC
TGTGGAGCAGAGGAATGAGATCCTGACCGCCATCCTGGCCTCCCTGACCGCCAGACAGAATCTGAGGAGAGAGTGGCACG
CCAGGTGTCAGTCCAGGATCGCCAGAACCCTGCCCGCCGATCAGAAGCCTGAGTGTAGACCCTACTGGGAGAAGGATGAT
GCCTCCATGCCCCTGCCCTTCGACCTGACAGACATCGTGAGCGAGCTGAGAGGCCAGCTGCTGGAGGCCAAGCCCGGCTC
TCACCACCACCACCATCACTGA
```

(2) Expression vector

The cDNA sequences described in §2.1 were subcloned in ProteoGenix's proprietary expression vector pATX21 (cf Figure S1). The expected proteins produced are illustrated below.

***> FTO-6His Protein- 513 AAs- 59.16kDa***

```
MKRTPTAEEREREAKKLRLLEELEDTWLPYLTPKDDEFYQQWQLKYPKLILREASSVSEELHKEVQEAFLTLHKHGCLFR
DLVRIQGKDLLTPVSRILIGNPGCTYKYLNTRLFTVPWPVKGSNIKHTEAEIAAACETFLKLNDYLQIETIQALEELAAK
EKANEDAVPLCMSADFPRVGMGSSYNGQDEVDIKSRAAYNVTLLNFMDPQKMPYLKEEPYFGMGKMAVSWHHDENLVDRS
AVAVYSYSCEGPEEESEDDSHLEGRDPDIWHVGFKISWDIETPGLAIPLHQGDCYFMLDDLNATHQHCVLAGSQPRFSST
HRVAECSTGTLDYILQRCQLALQNVCDDVDNDDVSLKSFEPAVLKQGEEIHNEVEFEWLRQFWFQGNRYRKCTDWWCQPM
AQLEALWKKMEGVTNAVLHEVKREGLPVEQRNEILTAILASLTARQNLRREWHARCQSRIARTLPADQKPECRPYWEKDD
ASMPLPFDLTDIVSELRGQLLEAKPGSHHHHHH
```

Feature: 6His tag with linker : [506: 513]

Figure S1. Map of the vector for ***FTO-6His Protein***

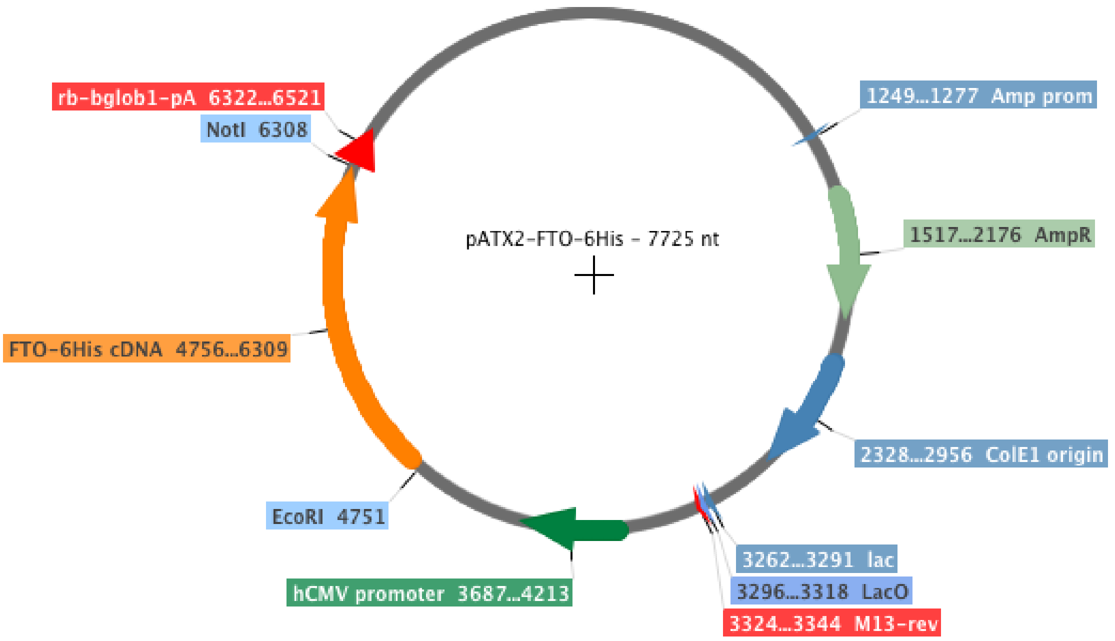


(3) Expression tests.

3.1 Protocol description

An endotoxin-free DNA preparation was done for each plasmid construction obtained as described in part 2. By using a proprietary method, the constructions were then transiently transfected in HEK293 cells. Culture medium and cells samples were collected at different time-points post-transfection in order to analyze target proteins expression.

3.2 Protein samples preparation and analysis

For each time-point, culture medium and cells samples were prepared and analyzed as followed. Culture medium samples were analyzed after semi-purification on nickel resin: Briefly, samples were incubated with the resin, and proteins bound on the resin were then analyzed by loading on SDS-PAGE. For cells samples, total protein extracts were prepared from cell lysates in native conditions (sonication in PBS pH 7.5) and denatured conditions (pellets after native lysis were solubilized in PBS pH 7.5 supplemented with urea). Native protein extracts (NPE) and denatured protein extracts (DPE) were then analyzed on SDS-PAGE after semi-purification conducted as described for culture medium samples.

3.3 The target protein was expressed at good levels in HEK293 cells. (Expression tests of the FTO are given on Figure S2).

Figure S2. **Expression test**. Coomassie blue staining. Reducing SDS-PAGE analysis of Medium, NPE and DPE samples prepared as described in p.3.2. Ø. Untransfected control culture. D3-D5. Days post-transfection.

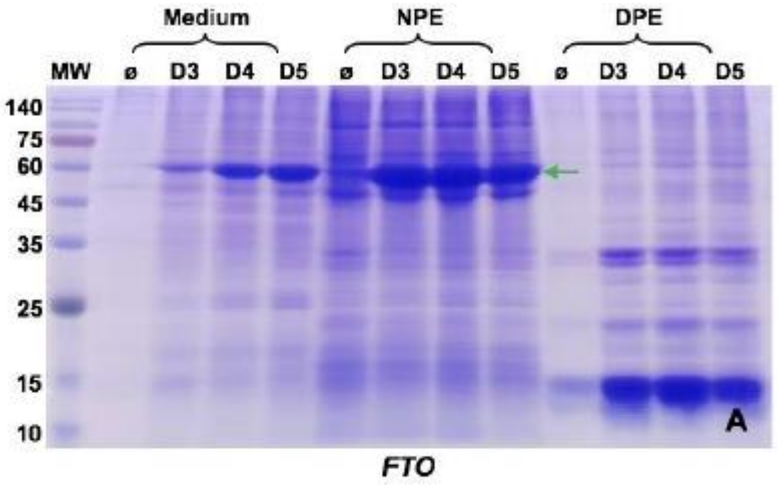


(4) Production and Purification Scale-ups

4.1 Purification short protocol description.

An 80ml-culture of HEK293 was transfected in optimal conditions described in Table1, in order to produce native/soluble target proteins. Affinity purification (IMAC) versus His-tag were then conducted by a standard method starting from NPE prepared as described in p. 3.2:

-Binding in PBS pH7.5

-Wash and elution by imidazole shift

-Analysis by SDS-PAGE of fractions of interest

-Final sample QC: qualitative by SDS-PAGE, quantitative by Bradford method

After purification, fractions of interest were pooled and buffer exchanged versus the following final buffer: PBS pH 7.3, 0.02% NaN3, 50%Glycerol.

4.2 Results.

The results of the FTO purification tests are given in Figure S3.

Figure S3. **FTO purification test**. Coomassie blue staining. Reducing SDS-PAGE analysis A. Purification profile. B. Final sample QC after buffer exchange (2µg). IN. Input sample. FT. Flow through. W1-W3. Wash steps. E1-E9. Eluted fractions. Molecular weight (MW) markers in kDa are shown in the left.

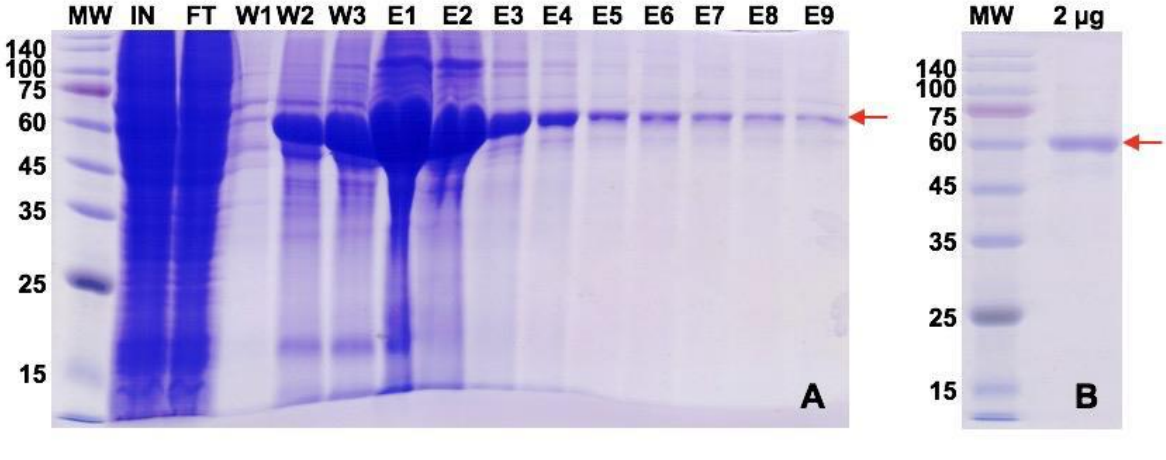


The results obtained for FTO are consistent with the data obtained for the small-scale tests. The purity of the protein was >90%.
